# Single-cell profiling reveals MMP7-associated epithelial transition and immune-stromal remodeling in esophageal carcinogenesis

**DOI:** 10.64898/2026.07.28.741095

**Authors:** Qi Wang, Peng Li, Li Fu, Yuewen Zhang, Wanli Zheng, Lin Li, Qinggang Hao, Wei Luo, Haiyan Guo, Mingyao Meng, Ruhong Li, Zongliu Hou, Yangfan Guo

## Abstract

The progression from reflux esophagitis (RE) to Barrett’s esophagus (BE) and esophageal adenocarcinoma (EAC) is a major route of inflammation-associated esophageal tumorigenesis, but the cellular transitions involved remain incompletely understood. We performed single-cell transcriptomic profiling of RE, BE, and EAC tissues and integrated public EAC datasets, yielding 51,283 cells across the RE-BE-EAC continuum. The analysis identified stage-associated epithelial and microenvironmental differences. Pseudotime analysis placed Basal epithelial cells along two differentiation branches, one toward mature epithelial states and the other through proliferative and gastric-type metaplastic states toward copy number alteration-bearing malignant epithelial cells. MMP7 increased along this malignant branch, was enriched in cancer-associated epithelial cells. Its knockdown reduced the proliferation and migration of OE19 and OE33 cells, supporting an association with proliferative and migratory phenotypes. In parallel, the microenvironment shifted toward immune suppression and stromal remodeling, with increased T-cell exhaustion, macrophage polarization toward tumor-associated states, and enrichment of CAF-like myofibroblasts. Ligand-receptor analysis further suggested an epithelial-centered communication network involving MDK-SDC2/LRP1/NCL interactions between malignant epithelial cells and fibroblast, T-cell, and myeloid compartments. These results identify MMP7-associated epithelial states and candidate MDK-related communication pathways for further longitudinal and functional validation.

## Background

Esophageal adenocarcinoma is the seventh most common cancer worldwide and the sixth leading cause of cancer-related mortality, characterized by high mortality rates, poor prognosis at diagnosis, and marked geographic variation in incidence^1,2^. In China, an estimated 224,000 new cases of esophageal cancer and 187,500 related deaths occurred in 2022, making it one of the leading causes of cancer mortality nationwide^3^. EAC represents a major histological subtype of esophageal cancer and is closely associated with gastroesophageal reflux disease and Barrett’s esophagus, a premalignant condition arising from chronic reflux-induced injury^4,5,6,7^. The American College of Gastroenterology guidelines identify BE as one of the most important precursor lesions for EAC development^8^. However, only a small proportion of patients with BE ultimately progress to high-grade dysplasia or EAC^9^, indicating that conventional histopathological classification alone is insufficient to accurately predict malignant progression risk. Therefore, the key issue is not merely to confirm the existence of the RE-BE-EAC disease sequence, but rather to identify the cellular states and molecular characteristics that truly change during inflammation, metaplasia, and malignant transformation, and that may indicate the risk of malignant progression.

Previous studies have explored the mechanisms underlying the progression from BE to EAC from multiple perspectives. For example, Nowicki-Osuch^11^ et al. demonstrated through single-cell transcriptomics, methylation lineage tracing, chromatin accessibility profiling, and organoid models that BE more closely resembles a metaplastic state derived from the gastric cardia and is regulated by transcriptional programs driven by c-MYC and HNF4A. Their findings further suggested that EAC may originate from an undifferentiated BE cell population and can even arise in the absence of histologically recognizable metaplastic precursor lesions^10^. Maag JLV et al. revealed widespread genomic instability during the transition from BE to EAC using whole-transcriptome sequencing, characterized by aberrant expression of key DNA repair genes (e.g., BRCA1 and PRKDC), dysregulation of long non-coding RNAs, and systematic downregulation of Alu elements^11^. Strasser et al. found that progression from BE to EAC is not merely an epithelial process. The replacement of metaplastic epithelium and alterations in tissue architecture are accompanied by concurrent changes in stromal cells, extracellular matrix composition, and tissue stiffness. Notably, fibroblasts exhibiting cancer-associated fibroblast (CAF)-like molecular features emerge during the premalignant metaplastic stage and express the immunosuppressive protein POSTN, indicating that BE progression is a multifaceted process involving coordinated interactions among epithelial, stromal, and immune compartments^12^. Collectively, these studies suggest that EAC development is not driven solely by genetic alterations or epithelial transformation, but rather results from the interplay between intrinsic epithelial reprogramming and continuous microenvironmental remodeling. However, most previous investigations have focused primarily on BE and EAC or on specific stages of disease progression, providing limited coverage of the continuous transition from reflux-induced inflammation to Barrett’s metaplasia and ultimately to EAC. In particular, during the RE-BE-EAC progression continuum, it remains unclear which epithelial cell states represent inflammatory repair responses, which reflect a shift toward malignant lineages, and how these epithelial states interact with other cellular components to establish a tumor-promoting microenvironment. These questions remain insufficiently addressed and warrant systematic investigation.

To address these knowledge gaps, we performed single-cell RNA sequencing on tissues from RE, BE and EAC, and integrated publicly available single-cell datasets of EAC. By constructing a single-cell atlas spanning inflammation, premalignant lesions and invasive carcinoma, we sought to characterize cellular heterogeneity, infer epithelial state trajectories, and compare immune and stromal compartments across diagnostic groups. Furthermore, we aimed to identify key cell-cell communication networks associated with the development of BE and the malignant transformation to EAC. This study provides a single-cell framework for investigating stage-associated cellular states in esophageal adenocarcinoma evolution and generates hypotheses for future biomarker and therapeutic studies.

## Methods

### 1. Patients

Single-cell transcriptomic and clinical data were collected from patients with RE, BE and EAC. A total of five samples, including two RE samples, two BE samples, and one EAC sample, were obtained from the Department of General Surgery *Ⅱ*, Yan’an Hospital Affiliated to Kunming Medical University. Public EAC scRNA-seq data from GEO (GSE273127) were integrated into the analysis, and an independent dataset (GSE292971) was used for validation. RE and BE tissues were collected from mucosal regions adjacent to the esophageal dentate line, whereas EAC tissues were obtained from tumors at the gastroesophageal junction. Written informed consent was obtained from all participants, and detailed clinical information is provided in **Table S1**.

### 2. Single-cell transcriptomic sequencing and data processing

Single-cell suspensions were stained with 0.4% trypan blue, counted, and adjusted to a concentration of 1,000-2,000 viable cells/μL. Single-cell libraries were generated using the 10× Genomics Chromium platform and sequenced on an Illumina platform with paired-end 150 bp reads.

Raw sequencing data were processed using Cell Ranger (v8.0.0, 10×Genomics) and aligned to the GRCh38 reference genome to generate gene expression matrices. Doublets were removed using DoubletFinder^13^, and batch effects were corrected using Harmony^14^. Downstream analyses were performed with the Seurat package^15^. Cells with fewer than 400 or more than 5,600 detected genes, or mitochondrial transcript proportions exceeding 20%, were excluded. Highly variable genes were identified for data normalization, principal component analysis (PCA), and Louvain clustering (top 10 PCs, resolution = 0.3, k = 30). UMAP was used for dimensionality reduction and visualization. Cell types were annotated based on SingleR predictions^16^, the CellMarker database, and canonical marker genes reported in previous studies^17^. Target cell populations were subsequently extracted for reclustering and subpopulation annotation according to their characteristic marker gene expression profiles.

### 3. Single-cell transcriptomic downstream analyses

Single-cell transcriptomic downstream analyses included copy number variation (CNV) inference, differential expression analysis, trajectory inference, differential abundance analysis, and virtual gene knockout analysis. CNVs were estimated using inferCNV^18^ with normal esophageal single-cell datasets as reference controls. Differentially expressed genes (DEGs) were identified using the FindMarkers and FindAllMarkers functions in Seurat (logFC > 0.25, expressed in ≥10% of cells), followed by Gene Ontology enrichment analysis using enrichGO^19^. Functional states of cell subsets were evaluated using GSVA^20^ and GSEA^21^. Cellular trajectories were reconstructed with Monocle3^22^ to infer state transitions, using marker-defined cell populations as the root state. Differential abundance analysis was performed using Milo^23^ to identify cell neighborhoods exhibiting significant abundance changes across conditions. Virtual gene knockout analysis was conducted using scTenifoldKnk^24^by comparing wild-type and simulated knockout gene regulatory networks, enabling identification of genes and pathways potentially regulated by the target gene.

### 4. Cell Culture and In Vitro Functional Assays

Human esophageal adenocarcinoma cell lines (OE19 and OE33) and the normal esophageal epithelial cell line HET-1A were purchased from Shanghai Fuheng Biotechnology and authenticated by short tandem repeat (STR) profiling. Cells were cultured in OE19-, OE33-, or HET-1A-specific media at 37°C in a humidified incubator with 5% CO₂. Lentiviral vectors carrying *MMP7*-targeting shRNA (PSC78842-1: CGTATCATATACTCGAGACTT, PSC78843-1:

CGTCATAGAAATAATGCAGAA) or negative control shRNA (sh-NC) were used for transfection. Cells were harvested 48 h after transfection for RT-qPCR, Western blotting, CCK-8, Transwell, and wound-healing assays.

Total RNA was extracted using TRIzol reagent and reverse-transcribed into cDNA using GoScript™ Reverse Transcription Mix, Oligo(dT) (Promega, Beijing, China) according to the manufacturer’s instructions. Quantitative real-time PCR (RT-qPCR) was performed using SsoFast EvaGreen Supermix (Bio-Rad, CA, USA). The primer sequences were as follows: *MMP7*-F:TGCAGTGATGTATCCAACCTATG: *MMP7*-R: TTGCTAAATGGAGTGGAGGAACA; β-actin-F : CCTTCCTGGGCATGGAGTC, β-actin-R:TGATCTTCATTGTGCTGGGTG.

Protein expression levels were analyzed by Western blotting^25^. Total proteins were separated by SDS-PAGE and transferred onto PVDF membranes (ISEQ00010, Millipore, Germany). After incubation with primary antibodies (1:1000) overnight at 4°C and HRP-conjugated secondary antibodies (1:5000), protein bands were visualized using an enhanced chemiluminescence (ECL) detection kit (PK10003, Proteintech, Wuhan, China).

Cell proliferation was evaluated using the Cell Counting Kit-8 (CCK-8) assay. Cells were seeded into 96-well plates and cultured for 24, 48, and 96 h. At each time point, cells were incubated with CCK-8 working solution for 2 h at 37°C, and absorbance was measured at 450 nm using a microplate reader. Cell migration was assessed using Transwell and wound-healing assays. For the Transwell assay, cells were seeded into the upper chamber, while complete medium containing 10% FBS was added to the lower chamber as a chemoattractant. After incubation for 16-18 h, migrated cells were fixed with methanol, stained with 0.1% crystal violet, and quantified using ImageJ software. For the wound-healing assay, confluent cell monolayers were scratched with a sterile 200 μL pipette tip and cultured in serum-free medium. Images were captured at 0 h and the indicated time points, and migration ability was evaluated by measuring wound closure rates.

### 5. T-cell and macrophage scoring analysis

T-cell functional states and activation levels were quantified using TcellSI^26^. Eight T-cell functional states, including quiescence, regulating, proliferation, helper, cytotoxicity, progenitor exhaustion, terminal exhaustion, and senescence, were evaluated from transcriptomic data. For each sample, expression values were first normalized using housekeeping genes. State-specific gene set expression ranks were then compared with theoretical distributions to calculate a T-cell state score (TCSS) ranging from 0 to 1, where higher scores indicate greater activity of the corresponding state. Macrophage functional signatures were evaluated using the AddModuleScore function in the Seurat package. M1- and M2-like macrophage scores were calculated based on marker gene sets derived from Azizi et al^27^.

### 6. Construction of a cell-cell interaction network

Cell–cell communication analysis was performed using CellChat^28^ to quantitatively infer potential interactions among different cell types, including ligand-receptor pairs and their cofactors. A CellChat object was constructed using the createCellChat function, and the Secreted Signaling database was selected for downstream analysis. Intercellular communication networks were inferred using the computeCommunProb, computeCommunProbPathway, and aggregateNet functions. Multiple CellChat objects were integrated using mergeCellChat. Finally, cell-cell interaction patterns were visualized using the netVisual_bubble function to illustrate intercellular crosstalk.

## Results

### 1. The single-cell transcriptomic landscape of reflux esophagitis, Barrett’s esophagus, and esophageal adenocarcinoma

In this study, tissue samples were collected from patients diagnosed with RE, BE and EAC. Ultimately, two RE samples, two BE samples, and one EAC sample were obtained and subjected to single-cell RNA sequencing analysis **(Figure 1A)**, together with publicly available single-cell data from esophageal adenocarcinoma^29^. We constructed single-cell transcriptomic landscapes of RE, BE, and EAC. After quality control and doublet removal, 4,459 doublets were identified and excluded from downstream analysis **(Figure S1A)**, resulting in a final dataset of 51,283 cells for further analysis. Principal component analysis based on Harmony-corrected embeddings revealed clear separation of samples along PC1 according to disease stage, consistent with disease progression from RE to BE to EAC. Biological replicates within RE and BE clustered closely, indicating effective batch correction, whereas EAC samples exhibited marked heterogeneity, reflecting the intrinsic diversity of tumor cells **(Figure S1B)**.

**Figure 1.**
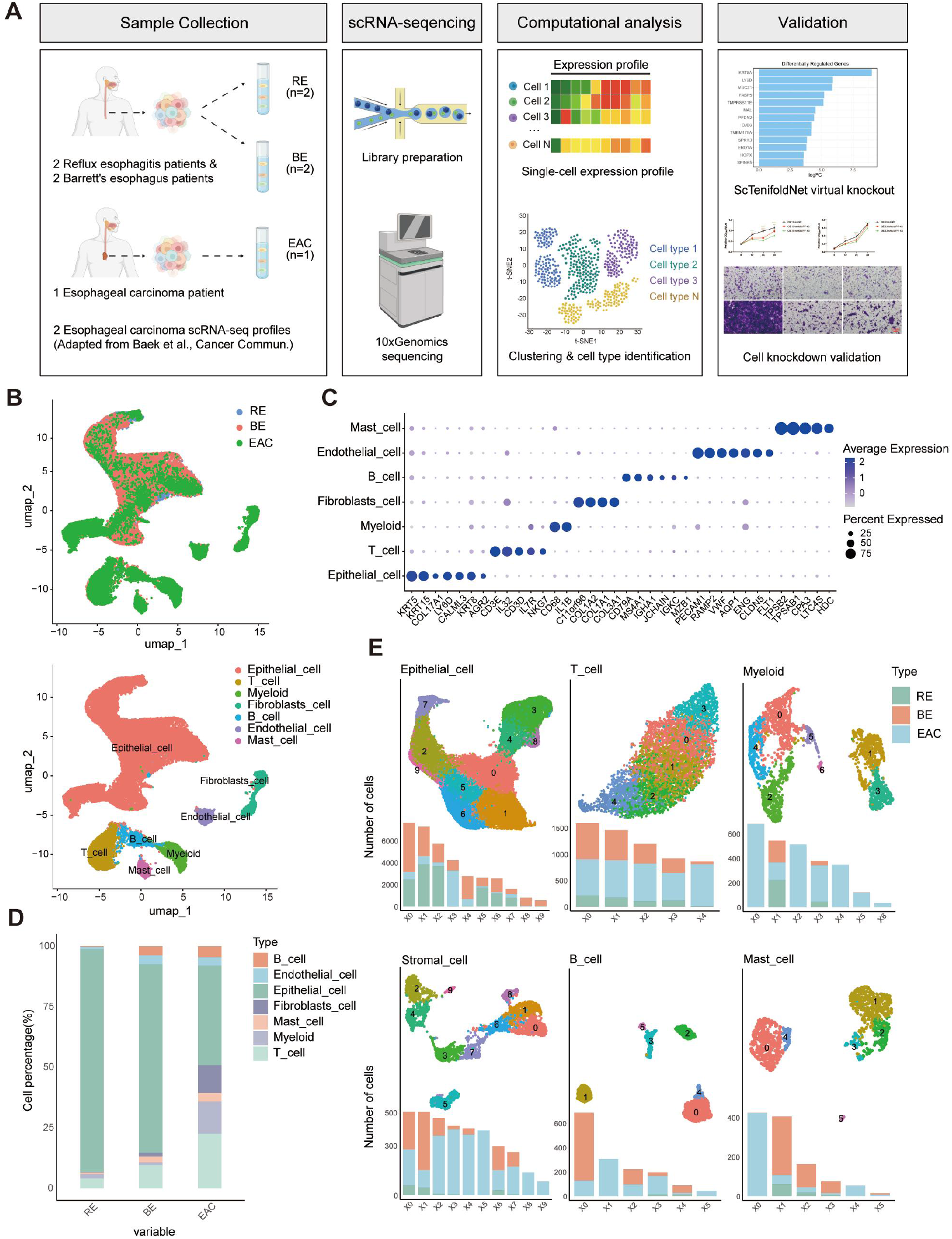
Overview of the ecological environment of RE, BE, and EAC as characterized by sc-RNAseq. a) Workflow for collecting and processing tissue samples of gastroesophageal reflux disease, Barrett’s esophagus, and esophageal cancer for scRNA-seq analysis. b) UMAP plot of 51283 cells, from top to bottom, respectively shows the transcriptomic features of disease types and sample cell types. c) Plot the point graph showing gene expression. d) The proportions of various cell types in samples of gastroesophageal reflux disease, Barrett’s esophagus, and esophageal cancer. e) UMAP plots and the number of subclusters for each major cell lineage, with these subclusters colored according to their respective sample sources.

Dimensionality reduction and unsupervised clustering were performed using the Seurat package to identify distinct cell populations. Based on UMAP visualization, cells selected for downstream analysis were classified into seven major clusters. According to canonical marker genes, these clusters were annotated as epithelial cells (*KRT5*, *KRT15*, *COL7A1*, *LY6D*, and *CALML3*), T cells (*CD3E*, *IL32*, *CD3D*, *IL7R*, and *NKG7*), myeloid cells (*IL1B* and *CD68*), fibroblasts (*COL1A2*, *COL3A1*, *C11orf96*, and *COL1A1*), B cells (*MS4A1*, *CD79A*, *JCHAIN*, *IGHA1*, *MZB1*, and *IGKC*), endothelial cells (*PECAM1*, *VWF*, *RAMP2*, *AQP1*, *CLDN5*, *ENG*, and *FLT1*), and mast cells (*TPSAB1*, *TPSB2*, *CPA3*, *HDC*, and *LTC4S*) **(Figure 1C)**. Comparative analysis of the proportions of these six major cell types across the three disease ecosystems revealed dynamic changes in cellular composition, highlighting the heterogeneity and complexity of the esophageal microenvironment **(Figure 1D)**. To further characterize cellular diversity, each major cell type was subjected to subclustering analysis to identify heterogeneous subpopulations. In total, 10 epithelial subclusters, 5 T-cell subclusters, 7 myeloid subclusters, 10 stromal subclusters, 6 B-cell subclusters, and 6 mast-cell subclusters were identified **(Figure 1E)**. Most subpopulations were shared among the three microenvironments, whereas several subsets were specifically enriched in esophageal adenocarcinoma, suggesting potential roles in the inflammation-to-cancer transition during esophageal tumorigenesis.

### 2. Putative Epithelial Trajectories and Functional Characterization of Progression-associated Genes Across RE, BE, and EAC

#### 2.1 The construction of the transcriptional landscape reveals the heterogeneity of epithelial cells

Considering the critical role of epithelial cells in esophageal cancer, we further characterized 35,853 epithelial cells derived from different esophageal lesions. Based on cluster-specific gene expression patterns and trajectory analysis, the 10 epithelial subclusters were annotated into distinct cell states. These included one basal epithelial cell cluster, Basal_Epi (*COL17A1*, *MIR205HG*, and *CD81*); three proliferating epithelial cell clusters, Prolif_Epi1, Prolif_Epi2, and Prolif_Epi3 (*LY6D*, *KRT5*, *SPRR1B*, and *TGM1*); two mature epithelial cell clusters, Mature_Epi1 and Mature_Epi2 (*SPRR3* and *SPRR1A*); three gastric-type metaplasia clusters, GTM1, GTM2, and GTM3 (*SMAD3*, *MUC6*, *PGC*, and *CLDN18*); and one putative malignant epithelial cell cluster, Malignant_Epi (*LGALS4*, *TSPAN8*, and *AGR2*) **(Figure 2A-B)**. Pseudotime analysis was used to infer putative state transitions among esophageal epithelial cells from basal epithelial cells to mature epithelial cells **(Figure 2C)**. Two putative branches were inferred: one branch differentiated from Basal_Epi to Mature_Epi, whereas the other progressed from Basal_Epi to Prolif_Epi and subsequently to GTM and Malignant_Epi. Based on marker gene expression and pseudotime analysis, Malignant_Epi was provisionally annotated as a malignant epithelial population. To validate their malignant features, copy number variation (CNV) analysis was performed by comparing Malignant_Epi cells with normal epithelial reference cells^30^. Compared with normal epithelial cells, Malignant_Epi cells exhibited widespread chromosomal amplifications and deletions, supporting their malignant characteristics **(Figure 2D)**. Furthermore, compositional analysis of epithelial subclusters across disease stages together with Milo analysis revealed a marked enrichment of Malignant_Epi cells in EAC relative to BE **(Figure 2E-F)**.

**Figure 2.**
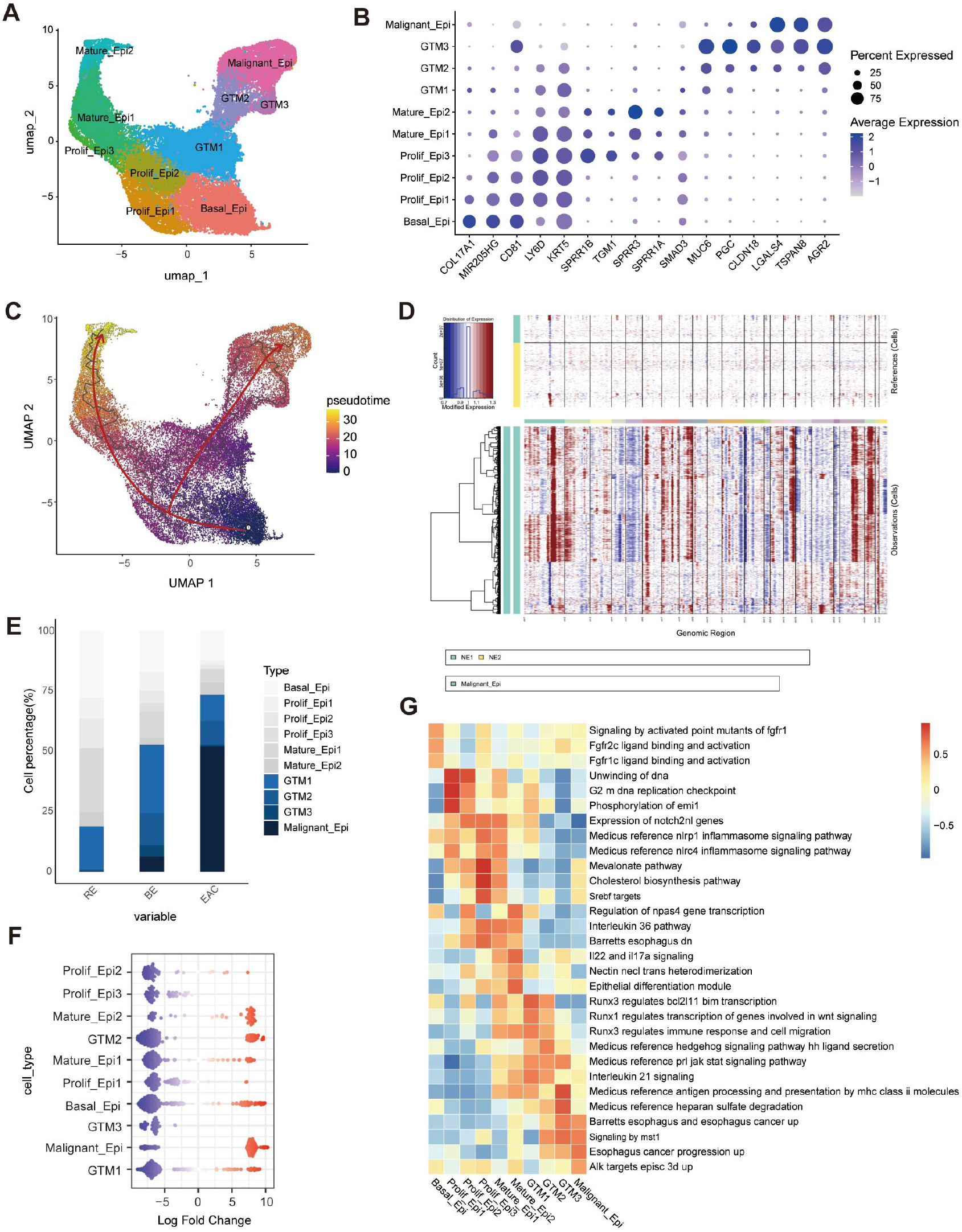
The heterogeneity of epithelial cells. a) UMAP of epithelial subclusters. b) Dot plot of marker gene expression. c) The pseudotime trajectory of epithelial cell subpopulations. d) CNV heatmap of epithelial cell clusters Malignant_Epi. Red: amplification, blue: deletion. The ranges of epithelial cells from different subclusters and different chromosomes are respectively marked by different color bars on the left and top of the heatmap. e) Cell proportions of each sub-cluster. f) Milo analysis of the difference in epithelial cell abundance between BE and EAC. The neighborhood abundance distribution of cell types is shown through the swarm plot. The blue and red dots represent decreased (logFC < 0) and increased (logFC > 0) cell abundance. g) The differences in pathway activity among the epithelial cell subpopulations scored by GSVA. Each column is standardized using z-score to represent the relative pathway activity.

To further investigate the heterogeneity among epithelial subclusters, GSVA was performed to evaluate pathway activities. Distinct pathway enrichment patterns were identified across different subclusters, whereas functionally related subtypes shared similar signaling features, highlighting the heterogeneity among epithelial cell populations **(Figure 2G)**. Notably, the MST1 and ALK signaling pathways were significantly enriched in the Malignant_Epi cluster. Dysregulation of MST1/2 signaling has been associated with tumor progression and immune dysfunction in multiple cancers, while ALK activation is a well-established oncogenic driver event. These findings further support the malignant characteristics of the Malignant_Epi cluster and suggest aberrant kinase signaling activation in these cells. Moreover, the activities of these pathways progressively increased along the differentiation trajectory from Basal_Epi to Prolif_Epi and subsequently to GTM and Malignant_Epi, indicating their potential involvement in the transition of esophageal epithelium from benign or precancerous states to malignant transformation.

#### 2.2 MMP7 is associated with the putative malignant epithelial trajectory and EAC cell proliferation and migration

Based on cell proportion and trajectory analyses, the Malignant_Epi cluster was predominantly derived from esophageal adenocarcinoma samples and exhibited highly malignant features. Therefore, subsequent analyses focused on the Malignant_Epi cluster. Pseudotime analysis demonstrated that the expression levels of Malignant_Epi marker genes, including *LGALS4*, *TSPAN8*, *AGR2*, and *MMP7*, progressively increased during disease progression **(Figure 3A and Figure S2A)**. To further validate these findings, an independent public dataset (GSE292971) was analyzed^31^ **(Figure S2B)**. Consistently, *LGALS4*, *TSPAN8*, and *AGR2* were expressed in both dysplastic and tumor samples, whereas MMP7, although expressed at a lower level, showed greater specificity in cancer samples. In addition, analysis of the TCGA database confirmed that *MMP7* was significantly upregulated in esophageal cancer **(Figure S2C)**. Therefore, *MMP7* was selected for further investigation. To investigate the regulatory role of MMP7, we performed an in silico knockout of *MMP7* in epithelial cells using scTenifoldKnk **(Figure 3B)**. *MMP7* depletion resulted in a set of significantly perturbed genes, indicating extensive rewiring of the underlying gene regulatory network. Gene set enrichment analysis of these perturbed genes revealed significant enrichment of esophageal secretory progenitor cell and late suprabasal epithelial cell signatures, suggesting that *MMP7* may contribute to the regulation of epithelial differentiation and maturation.

**Figure 3.**
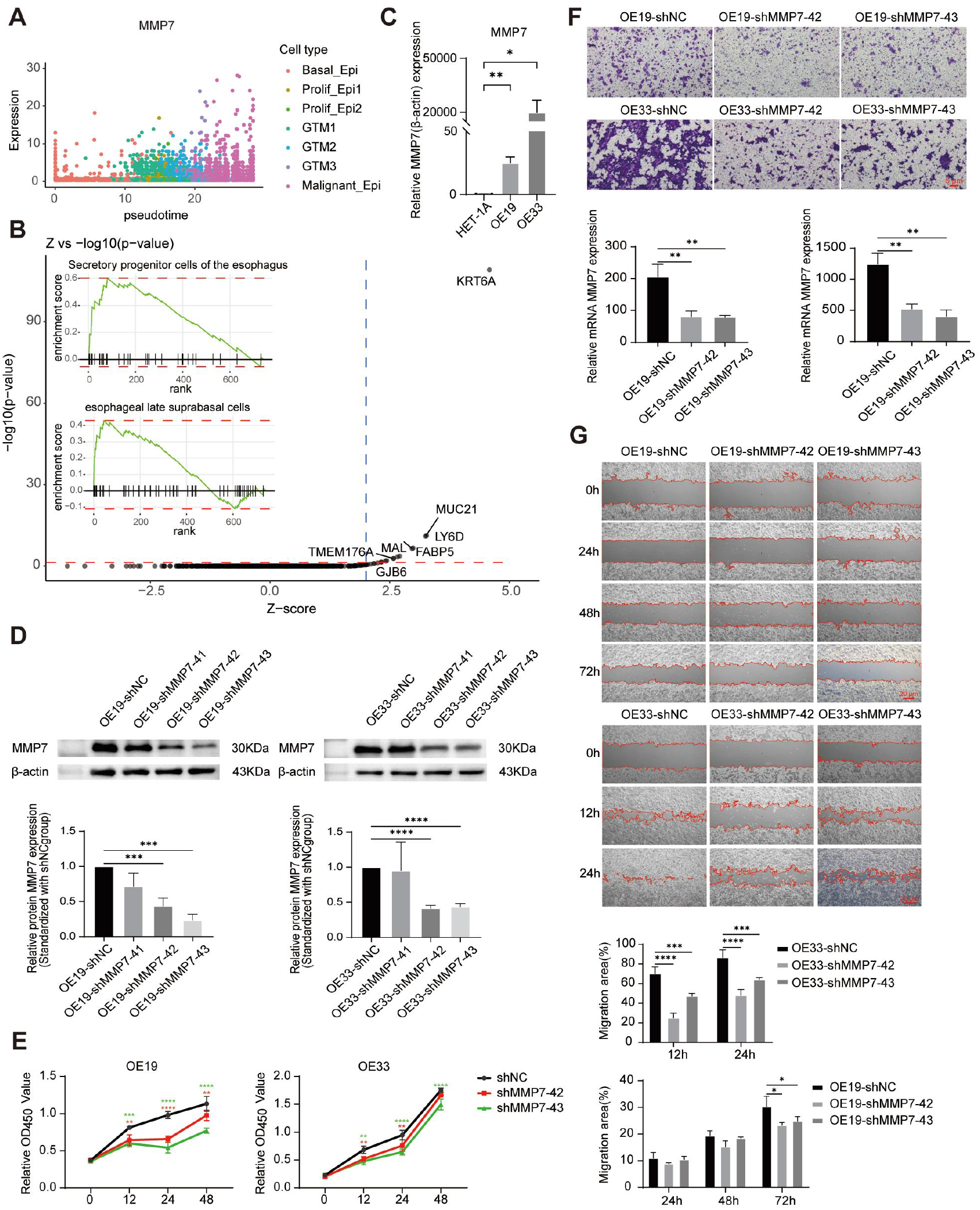
*MMP7* knockdown suppresses the proliferation and migration of esophageal cancer cells. a) The expression of the *MMP7* gene varies along the pseudo-time. b) The Q-Q plot of downstream genes after virtual knockout, with an inset showing the GSEA results of genes sorted by distance in the manifold-aligned scGRN. c) QRT-PCR was used to detect the expression level of *MMP7* in the HET-1A, OE19, and OE33 esophageal cell lines. d) Western blotting were used to verify th MMP7 knockdown in OE19 and OE33 cells. The data are presented as mean ± SD, * p < 0.05, * * p < 0.01, * * * p < 0.001. e-g) CCK-8, Transwell, and wound healing (scratch) assays were used to investigate the effects of *MMP7* knockdown on the proliferation and migration of esophageal cancer cells OE19 and OE33.

To further evaluate *MMP7* expression in esophageal adenocarcinoma cells, qRT-PCR was performed in normal esophageal epithelial cells and esophageal cancer cell lines. The results showed that *MMP7* expression was significantly elevated in the esophageal cancer cell lines OE19 and OE33 compared with the normal esophageal epithelial cell line HET-1A **(Figure 3C)**. Stable *MMP7*-knockdown cell lines were established using lentivirus-mediated shRNA transduction, and knockdown efficiency was validated by Western blot analysis. The results demonstrated that both shMMP7-42 and shMMP7-43 effectively reduced *MMP7* expression in the two esophageal cancer cell lines **(Figure 3D)**. To investigate the effect of MMP7 on esophageal cancer cell proliferation, CCK-8 assays were performed. Compared with the shNC control group, MMP7-knockdown cells exhibited consistently lower proliferation curves across multiple time points, and the differences became more pronounced with prolonged culture time, indicating that *MMP7* depletion significantly inhibited the proliferative capacity of esophageal cancer cells **(Figure 3E)**. In the Transwell assay, knockdown of *MMP7* significantly reduced the number of migrated OE33 cells on the lower membrane surface at 24h after seeding. In OE19 cells, a decrease in migrated cell numbers was observed at 48 h following seeding **(Figure 3F)**. In the wound-healing assay, *MMP7* knockdown significantly reduced the wound closure rate in OE33 cells at 24 h compared with the shNC control group, whereas a significant reduction was observed in OE19 cells at 72h **(Figure 3G)**. Taken together, these results indicate that *MMP7* is highly expressed in esophageal cancer, and its knockdown significantly suppresses cancer cell proliferation and migration. These findings suggest that *MMP7* may function as an oncogene in esophageal cancer, promoting tumor proliferation, migration, and progression.

### 3. Microenvironmental Remodeling During RE–BE–EAC Progression

#### 3.1 T-cell exhaustion-associated programs differ across RE, BE, and EAC

T cells represent a major component of the esophageal tumor microenvironment. To further investigate T-cell alterations during the inflammation-to-cancer transition, 6,032 T cells were reclustered and classified according to previously reported T-cell differentiation states^32^. Five T-cell subtypes were identified, including naïve T cells (*CCR7* and *MALAT1*), activated T cells (*RPS10* and *DUSP1*), effector T cells (expressing both activation- and cytotoxicity-related markers), cytotoxic T cells (*NKG7*, *CCL5*, *GZMB*, and *CD8A*), and exhausted T cells, which exhibited reduced cytotoxic marker expression together with increased expression of *TIGIT*, *CTLA4*, and *ENTPD1* **(Figure 4A-B)**. Pseudotime analysis placed naïve, activated, effector, and exhausted T-cell states along a putative continuum. **(Figure 4C)**. Quantitative analysis of T cells across different disease stages revealed that exhausted T cells were predominantly enriched in esophageal cancer samples, accompanied by reduced proportions of activated and effector T cells **(Figure 4D)**. Since the exhausted T cells (Effector) express high levels of CD39 (*ENTPD1*) and extremely low levels of *KLRG1*^33^, we hypothesize that the majority of the exhausted T cells are tumor-reactive T cells **(Figure 4E)**.

**Figure 4.**
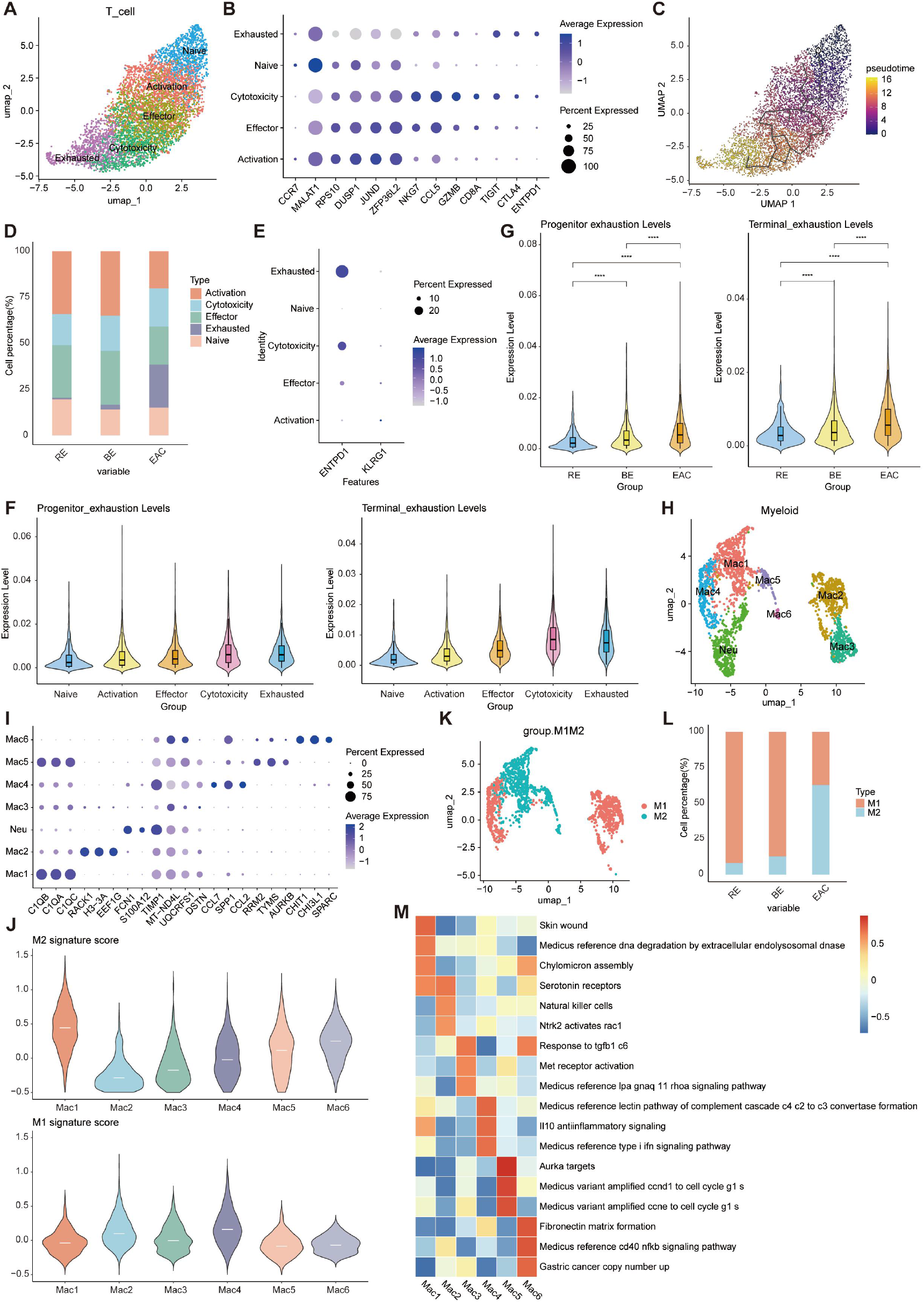
Changes in immune cells during the progression of esophageal cancer. a) UMAP plots of T cell subsets. b) Dot plots of marker gene expression. c) Pseudotime trajectories of T cell subsets. d) Proportions of cells in each T cell subset. e) Expression of ENTPD1 and KLRG1 genes in T cell subsets. f) Cytotoxicity and exhaustion scores calculated by the TcellSI package to quantify T cell characteristics. g) Initial and terminal exhaustion scores of exhausted T cells calculated by the TcellSI package. h) UMAP plot of myeloid cell subsets. i) Dot plot showing the expression of myeloid cell marker genes. j) Violin plots illustrating the pro-inflammatory (M1) and anti-inflammatory (M2) gene signatures of six macrophage clusters. White lines indicate the median scores for each gene signature. k) UMAP plot of pro-inflammatory (M1) and anti-inflammatory (M2) macrophage subsets. l) Proportions of pro-inflammatory (M1) and anti-inflammatory (M2) macrophages in esophageal samples across different disease types. m) Differences in pathway activity among macrophage subpopulations scored by GSVA. Each column is standardized by z-score to represent relative pathway activity.

To further characterize T-cell functional states, T-cell scores were calculated using the TcellSI package. Exhausted T cells showed the highest progenitor exhaustion and terminal exhaustion scores **(Figure 4F and Figure S3A)**. In addition, cytotoxic T cells also displayed relatively high terminal exhaustion scores, suggesting that exhaustion-associated programs may be activated concurrently with cytotoxic functions, a phenomenon commonly observed in tumors and chronic inflammatory conditions. Comparative analysis between cytotoxic and exhausted T-cell subsets revealed that cytotoxic T cells highly expressed effector molecules, including *GZMB*, *NKG7*, *PRF1*, and *GNLY*, whereas the expression of these cytotoxic genes was markedly reduced in exhausted T cells, which instead expressed inhibitory markers such as *TOX*, *CTLA4*, and *ENTPD1*. These findings indicate that the exhausted T-cell subset exhibited a more pronounced exhausted phenotype than the cytotoxic T-cell subset **(Figure S3B)**. Furthermore, progenitor exhaustion and terminal exhaustion scores were analyzed across disease stages, revealing a progressive increase in T-cell exhaustion during disease progression. Collectively, these cross-sectional data show higher exhaustion scores across the RE, BE, and EAC groups and are consistent with an increasingly immunosuppressive microenvironment **(Figure 4G)**.

#### 3.2 Macrophages subclusters exhibit stage-associated polarization programs

To investigate myeloid cell dynamics during inflammation-to-cancer transition, myeloid cells were reclustered and classified into six macrophage subclusters and one neutrophil cluster, including Mac1 (*C1QA*, *C1QB*, and *C1QC*), Mac2 (*RACK1*, *H3-3A*, and *EEF1G*), Mac3 (*DSTN*), Mac4 (*CCL7*, *SPP1*, and *CCL2*), Mac5 (*RRM2*, *TYMS*, and *AURKB*), Mac6 (*CHIT1*, *CHI3L1*, and *SPARC*), and Neu (*FCN1*, *S100A12*, and *TIMP1*) **(Figure 4H-I)**. Based on established M1/M2 and pro-/anti-inflammatory macrophage marker genes, macrophage polarization states were further evaluated^27,34^. Mac2 and Mac4 predominantly exhibited M1-like pro-inflammatory features, whereas Mac1, Mac5, and Mac6 displayed M2-like anti-inflammatory phenotypes. No clear polarization pattern was observed in Mac3 **(Figure 4J)**. Comparative analysis across disease stages revealed distinct macrophage polarization dynamics. Pro-inflammatory macrophages (Mac2 and Mac4) were enriched in RE and BE samples, whereas anti-inflammatory macrophages (Mac1, Mac5, and Mac6) were markedly increased in EAC samples **(Figure 4K-L)**. The relative abundance of macrophage subclusters shifted from M1-like populations in RE and BE toward M2-like populations in EAC, consistent with immune remodeling across diagnostic groups.

Further pathway analysis revealed distinct biological functions among macrophage subclusters. Pro-inflammatory macrophages (Mac2 and Mac4) showed enrichment of natural killer cell and IFN signaling pathways, both of which are key drivers of M1 polarization. In addition, activation of complement-related pathways, including C2, C3, and C4 signaling, further supported their pro-inflammatory phenotype. Anti-inflammatory macrophages displayed distinct functional characteristics. Mac1 was enriched in tissue repair and wound-healing pathways, consistent with classical M2 macrophage functions. Mac5 exhibited activation of E2F1 and E2F4 signaling pathways, together with upregulation of cell cycle regulators such as CCND1 and CCNE, indicating enhanced cell cycle activity and proliferative potential. Mac6 showed enrichment of a transcriptional gene set labeled gastric cancer copy number up; this enrichment does not by itself establish genomic alterations in macrophages **(Figure 4M)**.

#### 3.3 Tumor-associated changes in endothelial and myofibroblast subpopulation

To evaluate stromal cell dynamics within the tumor microenvironment (TME), stromal cells, including endothelial cells and fibroblasts, were extracted and reclustered. After subclustering and removal of contaminating cells, four endothelial cell subclusters were identified and annotated according to previously reported markers^35^, including two venous endothelial cell clusters, VenECs1 (*ACKR1*) and VenECs2 (*ACKR1* and *CXorf36*), one capillary-like endothelial cell cluster, CapECs (*RGCC*), and one lymphatic endothelial cell cluster, LECs (*CCL21* and *LYVE1*) **(Figure 5A-B)**. Cell proportion analysis and Milo analysis revealed that VenECs2 was specifically enriched in esophageal cancer, accompanied by a reduction in VenECs1, suggesting endothelial remodeling during tumor progression and indicating that VenECs2 may represent a tumor-associated venous endothelial subtype **(Figure 5C-D)**.

**Figure 5.**
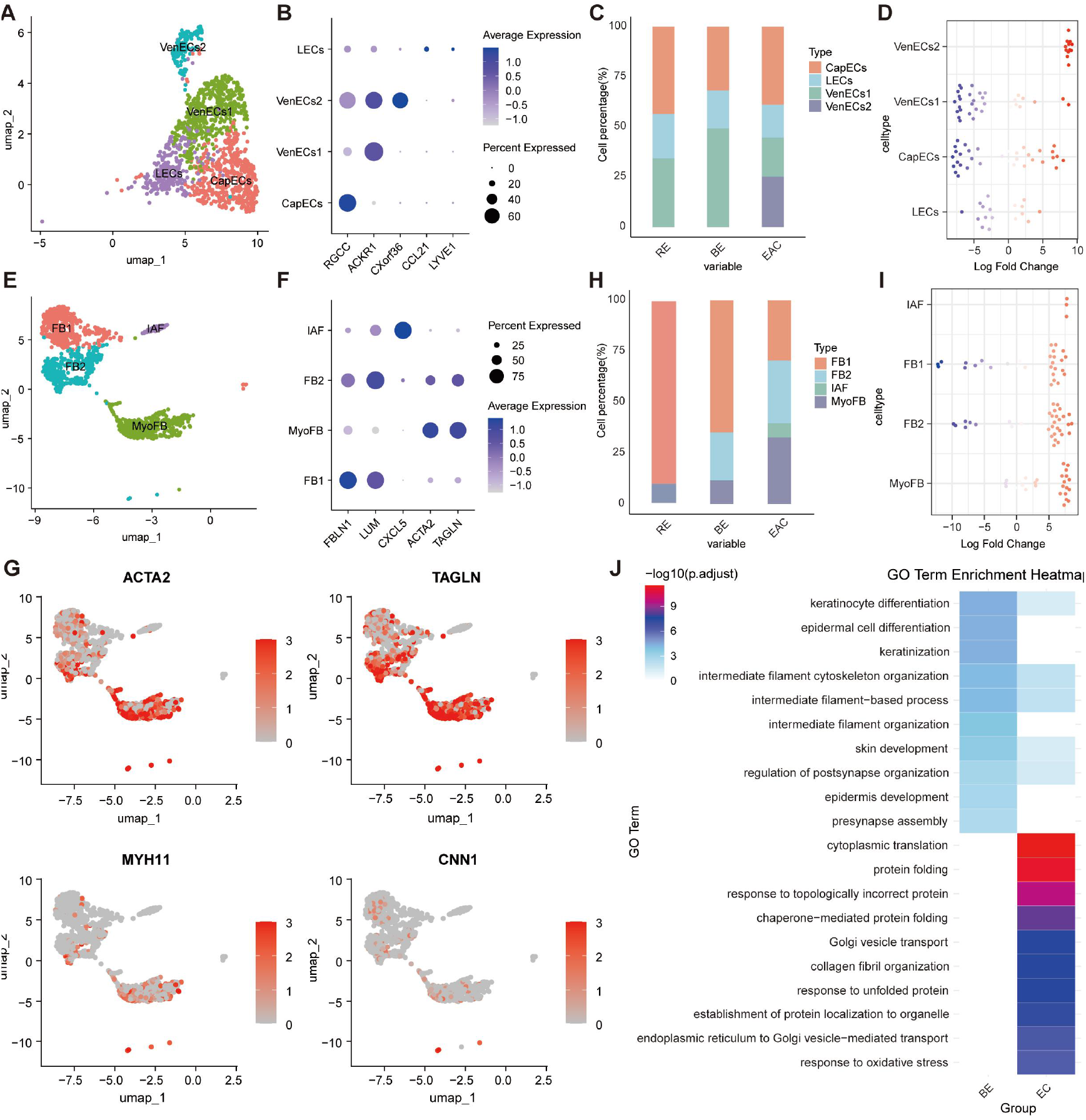
Changes in stromal cells during the progression of esophageal cancer. a) UMAP atlas of endothelial cell subpopulations. b) Dot plot of endothelial cell marker gene expression. c) Cell proportions of endothelial cell subpopulations. d) Milo analysis of differences in endothelial cell abundance between BE and EAC. e) UMAP atlas of fibroblast subpopulations. f) Dot plot of fibroblast marker gene expression. g) Cell proportions of fibroblast subpopulations. h) Milo analysis of differences in fibroblast abundance between BE and EC. i) Expression of ECM (extracellular matrix) genes, myofibroblast markers TAGLN and ACTA2, and smooth muscle cell genes MYH11 and CNN1 in fibroblast subpopulations. j) Gene Ontology (GO) analysis was conducted based on the differentially expressed genes of myofibroblasts in various disease types.

After subclustering and removal of contaminating cells, four fibroblast subclusters were identified and annotated according to previous studies^36^, including FB1 (*FBLN1*), FB2 (*LUM*), MyoFB (*ACTA2* and *TAGLN*), and IAF (*CXCL5*) **(Figure 5E–F)**. Previous studies have demonstrated that myofibroblasts play critical roles in tumor progression and stromal remodeling^37,38^. Because the MyoFB cluster highly expressed myofibroblast markers *ACTA2* and *TAGLN*, while exhibiting low expression of smooth muscle cell markers *MYH11* and *CNN1*, this cluster was defined as myofibroblasts **(Figure 5G)**. The cell ratio and Milo analysis revealed that MyoFB was significantly enriched in esophageal cancer **(Figure 5H-I)**. To further investigate MyoFB function, Gene Ontology (GO) analysis was performed using differentially expressed genes across disease stages. Compared with BE, MyoFBs in EAC were significantly enriched in pathways related to cytoplasmic translation, protein folding, and collagen fibril organization, suggesting functional reprogramming characterized by enhanced protein synthesis, extracellular matrix remodeling, and stress adaptation during tumor progression **(Figure 5J)**. Collectively, these findings indicate that myofibroblasts in esophageal cancer exhibit a typical cancer-associated fibroblast (CAF)-like activated phenotype.

### 4. The differences in intercellular communication within ecosystems of different disease types

To investigate how epithelial cells remodel the tumor microenvironment during disease progression, cell–cell communication analysis was performed across RE, BE, and EAC. Compared with RE, epithelial cell interactions with fibroblasts, B cells, and T cells were markedly increased in BE. In contrast, epithelial cell communication with fibroblasts and myeloid cells was significantly enhanced in EAC compared with BE **(Figure 6A-B)**. Ligand–receptor analysis further revealed progressive remodeling of epithelial cell-mediated signaling during disease progression. Notably, several pro-tumorigenic and immunoregulatory signaling axes, including MDK–SDC/LRP1, MIF–CXCR4/CD74, VEGF–VEGFR, and multiple laminin–integrin (LAMC/LAMB–ITGB1) interactions, were markedly enriched in EAC, suggesting enhanced epithelial-driven regulation of angiogenesis, immune cell recruitment, and stromal remodeling during tumor progression **(Figure 6C-D).** These interactions were further supported by ligand–receptor-specific expression patterns across epithelial–fibroblast, epithelial–T cell, and epithelial–myeloid cell communication networks **(Figure 6E)**. At the single-cell subtype level, the key epithelial-derived ligand MDK was upregulated in BE and further increased in malignant epithelial cells in EAC. Correspondingly, its cognate receptors (*SDC2*, *NCL*, and *LRP1*) were specifically expressed in cancer-associated fibroblasts (CAFs), exhausted T cells, and tumor-associated myeloid cells (Mac6), forming stage-specific and cell type-restricted communication axes. MDK interaction with *NCL*, *LRP1*, and *SDC2* has been reported to activate canonical pro-survival and proliferative pathways, including PI3K/Akt and MAPK/ERK signaling, thereby regulating cell proliferation, migration, adhesion, and inflammatory responses. Both *MDK* and *NCL* are frequently overexpressed in multiple malignancies and are associated with poor prognosis. Overall, these predictions suggest that epithelial signaling may differ across RE, BE, and EAC and may participate in stage-associated stromal and immune remodeling.

**Figure 6.**
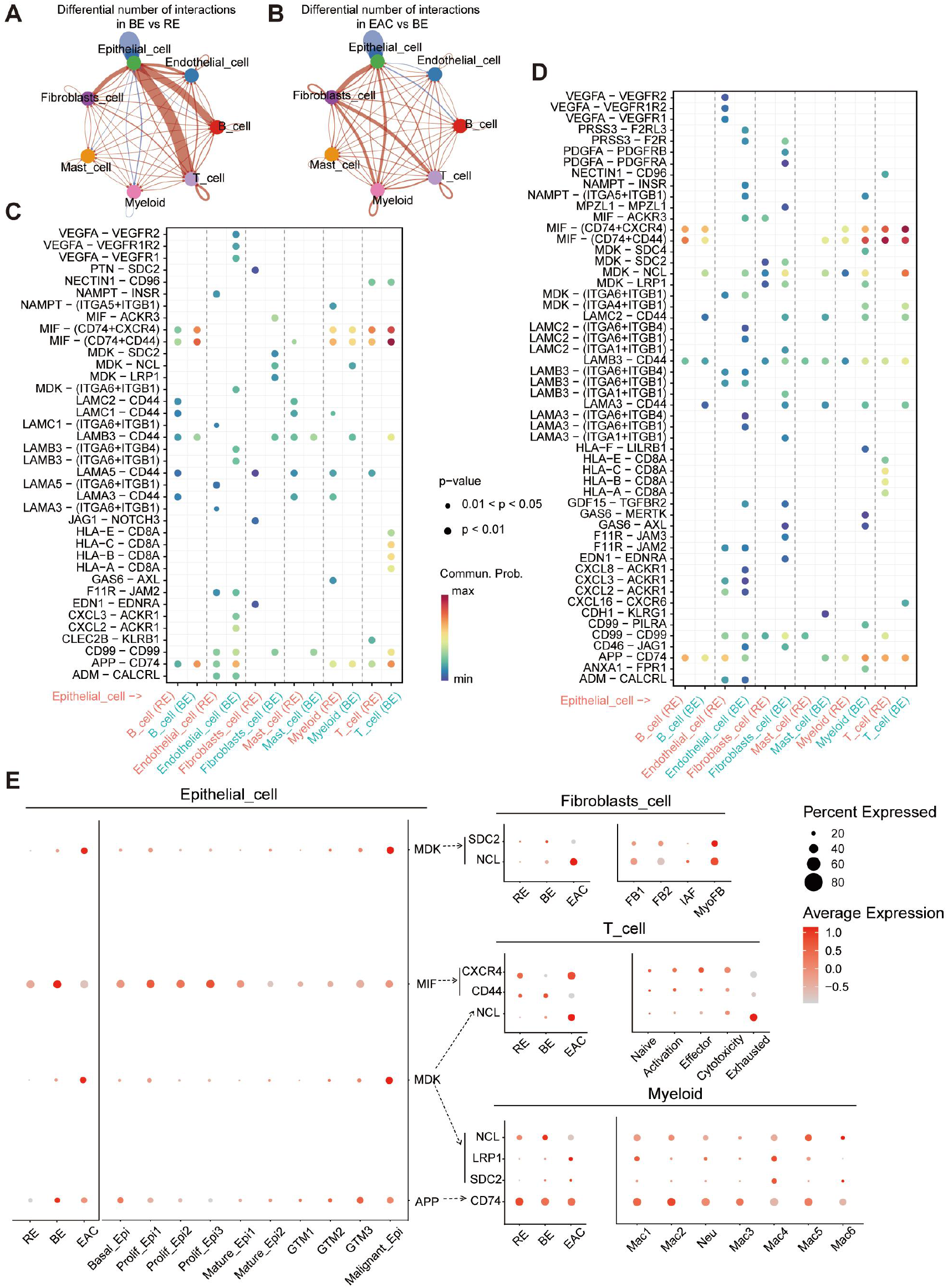
Differences in interactions and signals between various cell types has highlighted significant gene associations. a) A differential network map of the intercellular communication network between RE and BE cells, where red edges indicate increased signal transduction and blue edges indicate decreased signal transduction. b) A differential network map of the intercellular communication network between BE and EAC cells. c) A bubble chart showing the differences in all ligand-receptor pairs between RE and BE, with P-values represented by the size of the circles. d) A bubble chart showing the differences in all ligand-receptor pairs between BE and EAC. e) A dot plot of the gene expression of significantly upregulated signal transduction ligand-receptor pairs in the three disease types.

## Discussion

By integrating single-cell transcriptomic data from RE, BE and EAC, characterized cellular states associated with inflammation, premalignant lesions, and malignant tumors. Our analyses revealed that the transition from inflammation to malignancy is accompanied by coordinated alterations in epithelial cell states, immune functions, stromal remodeling, and intercellular communication. We identified epithelial subpopulations associated with malignant transformation and characterized their key molecular features, while also observed stage-associated differences in immune and stromal programs. Across RE, BE, and EAC, epithelial state differences coincided with T-cell exhaustion, shifts in myeloid programs, enrichment of CAF-like stromal populations, and predicted epithelial-immune-stromal communication **(Figure 7)**. Compared with previous studies that primarily focused on the BE-to-EAC transition, the inclusion of the RE stage in our analysis provides additional insight into distinguishing inflammation-associated reparative cell states from those associated with metaplastic transformation and malignant progression

**Figure 7.**
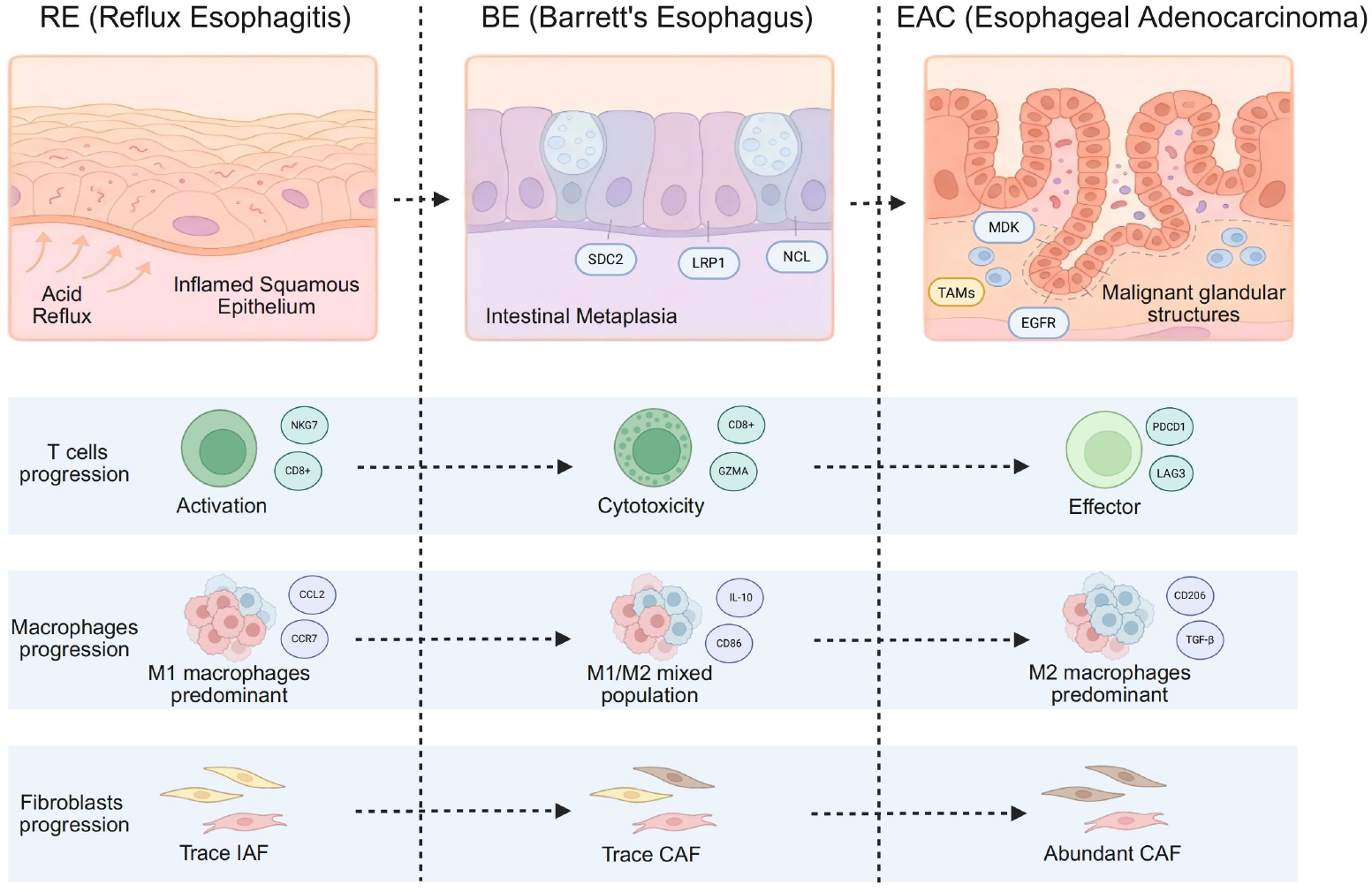
Progression from reflux esophagitis to esophageal adenocarcinoma.

At the epithelial level, pseudotime analysis inferred distinct putative trajectories from basal epithelial states. One trajectory terminated in a cluster annotated as malignant on the basis of marker expression, copy number alteration patterns, and enrichment in EAC; the analysis did not establish lineage ancestry or invasive behavior. These findings suggest that basal epithelial cells may not undergo malignant transformation through a simple linear process; rather, their cell fate may be redirected by microenvironmental cues, resulting in the emergence of distinct cellular states. However, the mechanism that controls this fate divergence remains unclear and requires further investigation. Furthermore, we identified *MMP7* as a gene markedly upregulated in malignant epithelial cells, a finding that was supported by external datasets and in vitro cell line experiments. Knockdown of *MMP7* significantly suppressed the proliferation and migration of OE19 and OE33 cells, suggesting that *MMP7* may promote the growth and migratory phenotypes of EAC cells. As a secreted matrix metalloproteinase, *MMP7* facilitates tumor cell migration and invasion through degradation of extracellular matrix components^39,40^. Combined with our trajectory analysis, these results raise the possibility that *MMP7* may function not only as a migration-associated effector molecule but also as a potential regulator of the transition toward malignant epithelial differentiation states. Longitudinal clinical validation would be required before *MMP7* could be evaluated as a marker of progression risk.

The immune microenvironment plays a critical role in cancer initiation and progression^41^. Pseudotime analysis placed naïve and exhausted T-cell states along an inferred continuum, and exhausted T cells were enriched in EAC. The relative abundance of macrophage subclusters shifted from predominantly pro-inflammatory programs in RE and BE toward anti-inflammatory programs in EAC, consistent with previous reports^42,43^. In addition, MyoFB cells were markedly enriched in EAC and displayed enhanced programs associated with extracellular matrix (ECM) remodeling, collagen organization, protein synthesis and secretion, and cellular stress responses. These findings suggest that CAF-like stromal activation may contribute to the establishment of the EAC microenvironment, in agreement with recent multi-omics studies proposing that BE-to-EAC progression is accompanied by coordinated alterations in both epithelial and stromal compartments^44,45,46^. Notably, stromal activation may further promote epithelial malignancy and immunosuppressive microenvironment formation by altering ECM composition, tissue stiffness, cell adhesion properties, and the spatial organization of immune cells. Nevertheless, the absence of spatial transcriptomic data in the present study precludes determination of whether MyoFB cells are preferentially localized around GTM, malignant epithelial cells, or specific immune cell populations to form distinct local microenvironments. Future studies incorporating spatially resolved approaches will be necessary to clarify the spatial architecture and functional interactions of these stromal cell populations during EAC progression.

Cell-cell communication analysis further suggested that epithelial cells progressively strengthened their potential interactions with stromal and immune cells during the transition from BE to EAC. Among these, the MDK-SDC2/LRP1/NCL signaling axis may serve as a putative epithelial-derived communication hub linking malignant epithelial cells, CAF-like fibroblasts, exhausted T cells, and tumor-associated myeloid cells. In addition, MIF-CXCR4/CD74 and VEGF-VEGFR interactions indicated coordinated remodeling of immune regulation, angiogenesis, and ECM-associated adhesion and migration programs. These findings suggest that epithelial cells undergo malignant transformation through intrinsic transcriptional reprogramming. Beyond these cell-autonomous changes, predicted ligand-receptor interactions suggest that epithelial cells may participate in broader remodeling of the tumor microenvironment. However, it should be noted that CellChat analysis infers potential cellular interactions solely on the basis of ligand and receptor expression and therefore cannot demonstrate spatial proximity, direct protein-protein interactions, or downstream signaling activation. Spatial co-localization analyses, protein-level assays, and MDK/receptor blockade experiments are needed to further validate the functional relevance of these predicted communication networks.

Several limitations of this study should be acknowledged. First, the disease trajectory was reconstructed using cross-sectional samples obtained from different patients. Substantial inter-individual differences in genetic background, immune status, microbiome composition, lifestyle, and medication history represent unavoidable confounding factors. Consequently, the observed differences in cellular composition and gene expression may reflect not only biological changes associated with disease progression but also patient-specific variation. Moreover, although RE, BE, and EAC represent relatively well-defined clinical diagnoses, the transition from inflammation to metaplasia and ultimately to malignancy constitutes a molecular continuum. Therefore, our sampling strategy may have overlooked important intermediate or transitional states during disease progression^47^. Second, because single-cell RNA sequencing requires tissue dissociation, spatial information is lost, limiting the ability to characterize the spatial organization of cellular populations and their interactions within the tissue microenvironment. In addition, the key cellular subsets, regulatory factors, and cell-cell communication networks identified in this study were primarily inferred from bioinformatic analyses and in vitro validation, and lack *in vivo* functional validation and confirmation in large independent cohorts. As a result, their clinical applicability remains to be fully established. Future studies integrating longitudinal sampling, spatial omics technologies, multi-omics approaches, and functional experiments will be essential for further elucidating the mechanisms underlying RE-BE-EAC progression and facilitating the clinical translation of these findings.

## Conclusions

In conclusion, this study characterizes stage-associated epithelial, immune, and stromal states across RE, BE, and EAC at single-cell resolution. The analyses identify a putative malignant epithelial trajectory marked by increasing *MMP7* expression, while cell-line experiments associate *MMP7* with proliferation and migration. Immune and stromal changes coincided with predicted epithelial-centered communication networks. These findings provide hypotheses for longitudinal, spatial, and functional validation rather than direct evidence of biomarkers or therapeutic targets.

## List of abbreviations

Basal_Epi: Basal Epithelial cells
BE: Barrett’s Esophagus
CAF: Cancer-Associated Fibroblast
CapECs: Capillary-like Endothelial cells
EAC: Esophageal Adenocarcinoma
ECM: Extracellular Matrix
EMT: Epithelial Mesenchymal Transition
FB: Fibroblast
GTM: Gastric Type Metaplasia
IAF: Inflammation-Associated Fibroblast
LECs: Lymphatic Endothelial cells
Mac: Macrophage
Malignant_Epi: Malignant Epithelial cells
Mature_Epi: Mature Epithelial cells
MyoFB: Myofibroblast
Prolif_Epi: Proliferating Epithelial cells
RE: Reflux Esophagitis
TME: Tumor Microenvironment
VenECs: Vascular Endothelial cells

## Declarations

### Ethics approval and consent to participate

This study received ethical approval from the Medical Ethics Committee of Yan’an Hospital Affiliated to Kunming Medical University. Informed consent was obtained from all participants enrolled in the study. (Number: 2023-082-01).

### Consent for publication

Not applicable.

### Availability of data and materials

The data reported in this paper have been deposited in the OMIX, China National Center for Bioinformation/Beijing Institute of Genomics, Chinese Academy of Sciences (https://ngdc.cncb.ac.cn/omix/release/OMIX018259). The public datasets analyzed in this study are available from the Gene Expression Omnibus (GEO) database under the accession numbers GSE273127, GSE292971, and GSE197677.

### Authors’ contributions

Qi Wang, Peng Li, Ruhong Li and Yangfan Guo conceived and designed the study. Zongliu Hou and Yangfan Guo are co-corresponding authors; they supervised the project and were responsible for project administration. Qi Wang collected and analyzed the data, and wrote the original draft of the manuscript. Peng Li and Yuewen Zhang provided clinical samples and resources. Wanli Zheng and Qinggang Hao was involved in data analysis. Wei Luo, Lin Li, Mingyao Meng and Haiyan Guo contributed to the writing, review, and editing of the manuscript. Li Fu performed the cell function experiments. All authors have read and approved the final manuscript for publication.

## Acknowledgements

The authors thank the clinical and laboratory staff involved in patient recruitment, sample collection, and data management for their support.Authors’ information (optional)

