## Supplementary material for "Single-cell profiling reveals MMP7-associated epithelial transition and immune-stromal remodeling in esophageal carcinogenesis"

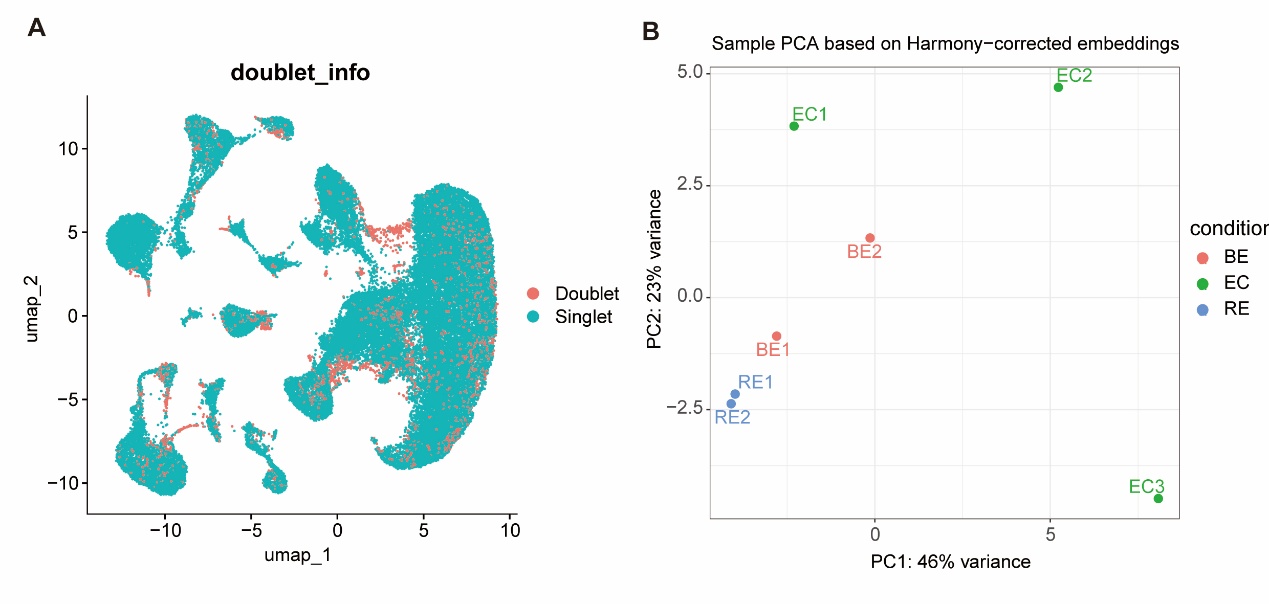

**Supplementary Figure1. The quality control status of the samples.**

a) UMAP plot showing the distribution of doublets.

b) Principal component analysis (PCA) plot after Harmony batch correction. Biological replicates within RE and BE clusters are tightly grouped, indicating effective batch correction, whereas EAC samples exhibit pronounced heterogeneity, reflecting the intrinsic diversity of the tumor.

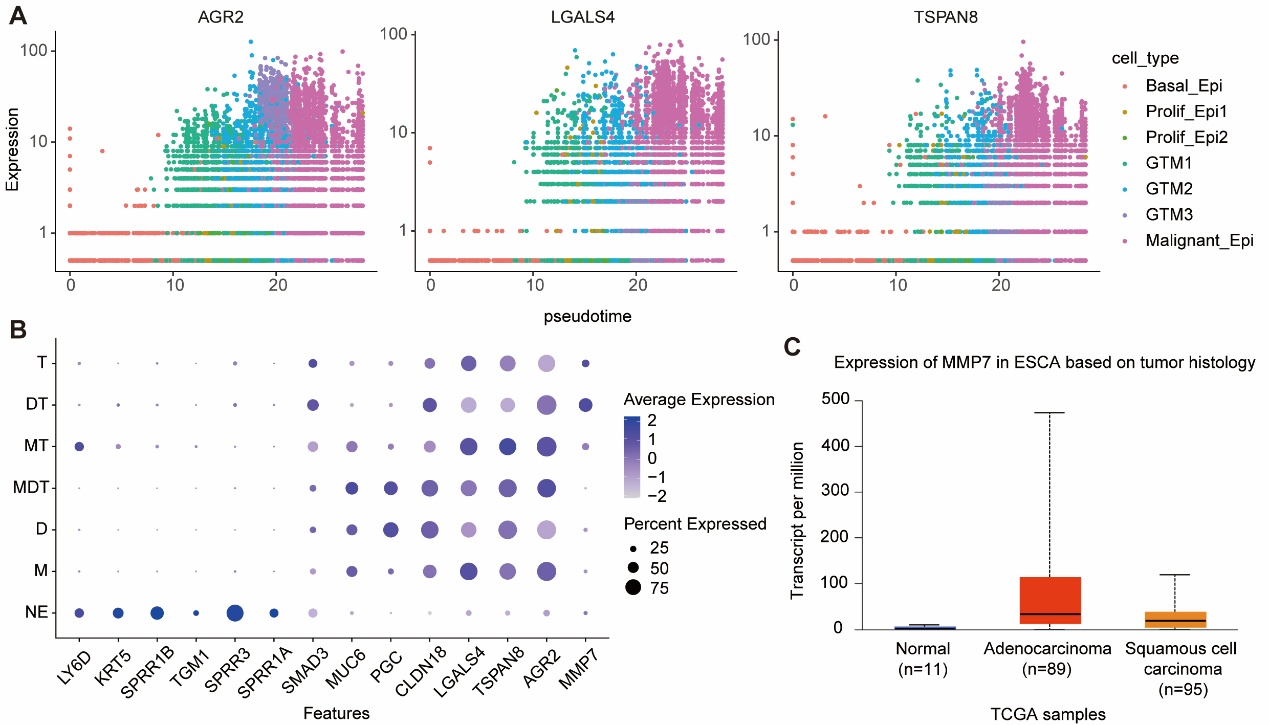

**Supplementary Figure2. The expression status of specific genes in the public database.**

a) Expression dynamics of AGR2, LGLAS4, and TSPAN8 along pseudotime.

b) Expression levels of AGR2, LGLAS4, TSPAN8, and MMP7 in the public database (GSE292971). NE: normal esophagus; M: metaplasia; D: dysplasia; T: esophageal adenocarcinoma.

c) Expression of MMP7 in esophageal cancer data from the TCGA database.

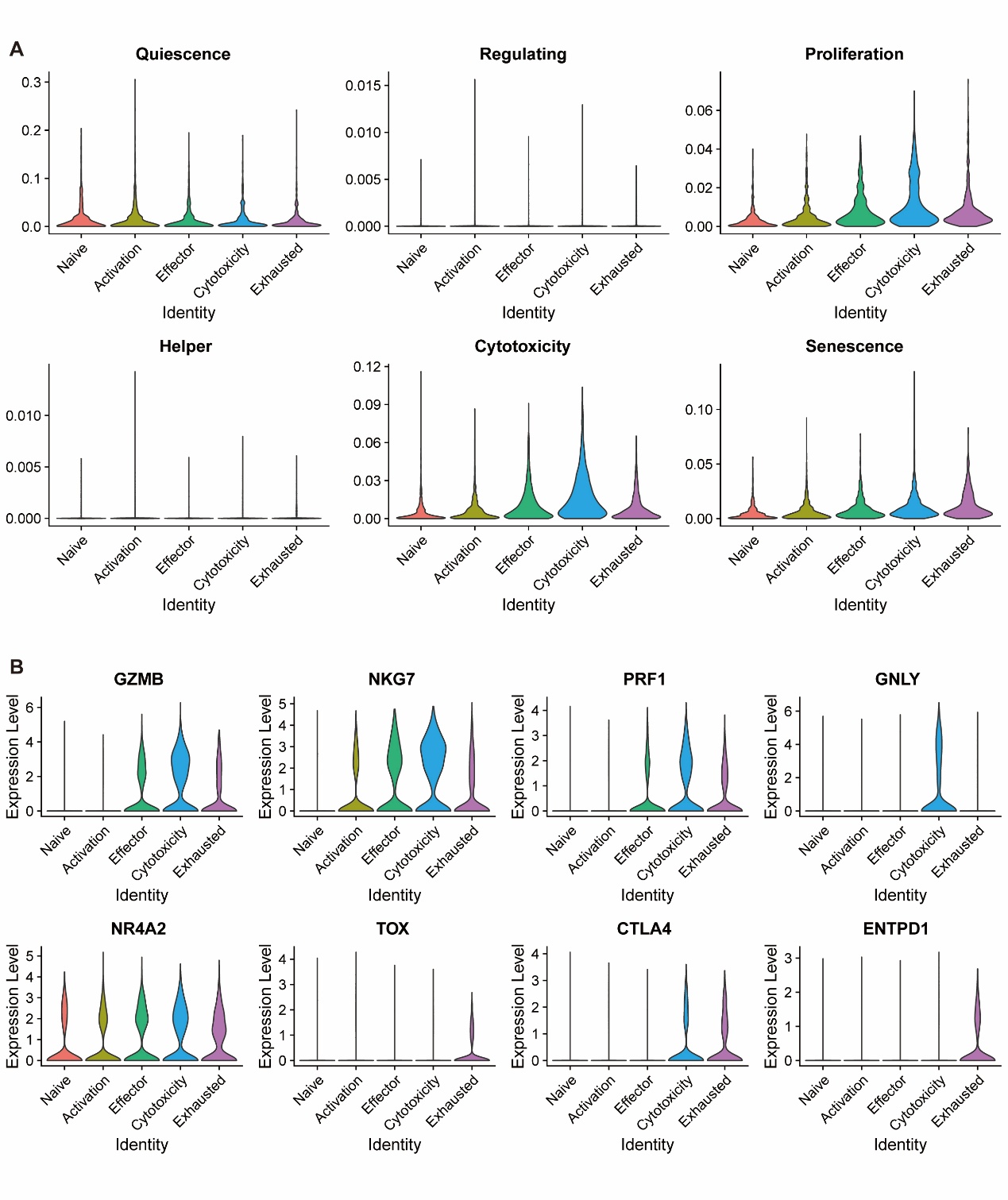

**Supplementary Figure3.** **T-cell subtype TcellSI score.**

a) Assessment of cell states across various cell types using TCellSI.

b) Expression of cytotoxicity-related genes and inhibitory genes across different cell types.

**Supplementary Table1 Clinical information of enrolled patients**

| **Sample ID** | **Age** | **Gender** | **Sample type** | **Tumor differentiation** |
| --- | --- | --- | --- | --- |
| RE-1 | 46 | M | RE | - |
| RE-2 | 65 | F | RE | - |
| BE-1 | 45 | M | BE | - |
| BE-2 | 69 | F | BE | - |
| EAC-1 | 60 | M | EAC | IIIA |
| EAC-2 | 71 | M | EAC | IIIB |
| EAC-3 | 44 | M | EAC | IIIB |
